# Locomotion enhances neural encoding of visual stimuli in mouse V1

**DOI:** 10.1101/102673

**Authors:** Maria C. Dadarlat, Michael P. Stryker

## Abstract

Neurons in mouse primary visual cortex (V1) are selective for particular properties of visual stimuli. Locomotion causes a change in cortical state that leaves their selectivity unchanged but strengthens their responses. Both locomotion and the change in cortical state are initiated by projections from the mesencephalic locomotor region (MLR), the latter through a disinhibitory circuit in V1. The function served by this change in cortical state is unknown. By recording simultaneously from a large number of single neurons in alert mice viewing moving gratings, we investigated the relationship between locomotion and the information contained within the neural population. We found that locomotion improved encoding of visual stimuli in V1 by two mechanisms. First, locomotion-induced increases in firing rates enhanced the mutual information between visual stimuli and single neuron responses over a fixed window of time. Second, stimulus discriminability was improved, even for fixed population firing rates, because of a decrease in noise correlations across the population during locomotion. These two mechanisms contributed differently to improvements in discriminability across cortical layers, with changes in firing rates most important in the upper layers and changes in noise correlations most important in layer V. Together, these changes resulted in a three- to five-fold reduction in the time needed to precisely encode grating direction and orientation. These results support the hypothesis that cortical state shifts during locomotion to accommodate an increased load on the visual system when mice are moving.

**Significance Statement:** This paper contains three novel findings about the representation of information in neurons within the primary visual cortex of the mouse. First, we show that locomotion reduces by at least a factor of three the time needed for information to accumulate in the visual cortex that allows the distinction of different visual stimuli. Second, we show that the effect of locomotion is to increase information in cells of all layers of the visual cortex. Third we show that the means by which information is enhanced by locomotion differs between the upper layers, where the major effect is the increasing of firing rates, and in layer V, where the major effect is the reduction in noise correlations.

## 1 Introduction

Behaviors such as locomotion, attention, and arousal have been shown to modulate cortical state (Niell and Stryker, 2010; Harris and Thiele, 2011; Ayaz et al., 2013; Bennett et al., 2013; Polack et al., 2013; Erisken et al., 2014; Reimer et al., 2014; Vinck et al., 2015). Locomotion, for example, increases stimulus-evoked neural firing in primary visual cortex of mice (Niell and Stryker, 2010) and possibly in the lateral geniculate nucleus (Niell and Stryker, 2010; Erisken et al., 2014). In mouse V1, the increase in firing rates is thought to be produced by disinhibiting pyramidal cells via a circuit separate from that which conveys visual input to V1, but see (Polack et al., 2013; Pakan et al., 2016; Dipoppa et al., 2016). Locomotion can be elicited via descending projections from the mesencephalic locomotor region (MLR) (Shik et al., 1966), which also send ascending projections to excite neurons in the basal forebrain (Nauta and Keypers, 1964), from which cholinergic projections to V1 activate a specific disinhibitory circuit (Fu et al., 2014; Pfeffer et al., 2013; Lee et al., 2014; Reimer et al., 2014). Importantly, stimulation of MLR can drive a change in cortical state even in the absence of overt locomotion (Lee et al., 2014), and could thus coordinate the initiation of locomotion and the change of cortical state.

What are the ethological and computational functions of this coordination? We hypothesize that the purpose of coupling cortical-state modulation with locomotion is to increase visually-relevant information encoded in the V1 neural population during periods in which visual information is expected to rapidly change, such as during locomotion. Consistent with this hypothesis, locomotion not only increases single-neuron firing rates but also, via heightened arousal, decorrelates neural spiking (Erisken et al., 2014; Vinck et al., 2015), both of which may contribute to increasing information within V1. Studies of single cells and mouse behavior further support this hypothesis: locomotion increases the rate with which mice detect low-contrast stimuli (Bennett et al., 2013) and depolarizes neural membrane voltages while decreasing their variability (Pinto et al., 2013; Polack et al., 2013).

These results strengthen our expectation that locomotion should increase the information content of V1 activity. However, increasing information in single neurons during electrical stimulation or behavior does not ensure that locomotion will increase information in the population of V1 neurons. For example, spontaneous transitions from low to high population firing rates in monkey V1 only shift information content among cells and do not increase information (Arandia-Romero et al., 2016).

Here, we use high-density microelectrode recording to test the hypothesis that populations of neurons in mouse V1 contain more information about visual stimuli during locomotion by decoding the direction and orientation of drifting gratings from single-trial population responses in different behavioral conditions. We find that locomotion does increase the information content of a neural population by at least two mechanisms, raising the firing rates of individual neurons and reducing noise correlations between neurons. These two mechanisms act cooperatively, and are present across all cortical layers, though to different extents. Increasing neural firing rates enhanced the information content of individual neurons and improved visual stimulus discriminability in population responses. Furthermore, even for trials with the same population firing rate, a decrease in pairwise noise correlations during locomotion further differentiated the representation of different gratings by V1. Together, these results suggest a computational function for the locomotion-induced modulation of neural firing and explain how this function is implemented. Our findings are consistent with a recent report that used 2-photon calcium imaging to study the responses of upper-layer excitatory neurons, which found increased information about the orientation of grating stimuli during locomotion as a result of increased firing rates, in particular for stimuli with high spatial resolution (Mineault et al., 2016).

## 2 Methods

### Animal procedures

Experiments were performed on adult C57/B16 mice (age 2-6 months) of either sex. The animals were maintained in the animal facility at the University of California, San Francisco (UCSF) and used in accordance with protocols approved by the UCSF Institutional Animal Care and Use Committee. Animals were maintained on a 12 hr light/12 hr dark cycle. Experiments in four mice were performed during the light phase of the cycle, and in four mice were performed during the dark phase of the cycle. We found no consistent differences in the results obtained from recording in animals in either light phase, and so pooled results from all animals.

### Preparation of mice for extracellular recording on the spherical treadmill

Our spherical treadmill was modified from the design described in (Niell and Stryker, 2010). Briefly, a polystyrene ball formed of two hollow 200 mm diameter halves (Graham Sweet Studios) was placed on a shallow polystryene bowl (250 mm in diameter, 25 mm thick) with a single air inlet at the bottom. Two optical USB mice, placed 1 mm away from the edge of the ball, were used to sense rotation of the floating ball and transmitted signals to our data analysis system using custom driver software.

During experiments, the animals head was fixed in place by a steel headplate that was screwed into a rigid crossbar above the floating ball. The headplate, comprised of two side bars and a circular center with a 5 mm central opening, was cemented to the skull a week before recording using surgical procedures as described in (Niell and Stryker, 2010). Briefly, animals were anesthetized with isoflurane in oxygen (3% induction, 1.5% maintenance) and given a subcutaneous injection of carprofen (5 mg/kg) as a postoperative analgesic, and a subcutaneous injection of 0.2 mL of saline to prevent postoperative dehydration. After a scalp incision, the fascia was cleared from the surface of the skull and a thin layer of cyanoacrylate (Vet-Bond, WPI) was applied to provide a substrate to which the dental acrylic could adhere. The metal headplate was then attached with dental acrylic, covering the entire skull except for the region in the center of the headplate, which was covered with a 0.2% Benzethonium chloride solution (New-Skin Liquid Bandage) to protect the skull. The animal was then allowed to recover. Three to seven days following headplate attachment, the animal was allowed to habituate to the recording setup by spending progressively more time on the floating ball over the course of two to three days (15 minutes to 1 hour), during which time the animal was allowed to run freely on the floating ball.

### Extracellular Recording in Awake Mice

The recording was performed as described previously (Niell and Stryker, 2010) with little modification. On the day of recording, the animal was again anesthetized as described above. The liquid bandage was removed, and the skull was thinned and removed to produce a craniotomy approximately 1-2 mm in diameter above the monocular zone of V1 (2.5 - 3 mm lateral to lambda). This small opening was enough to allow insertion of a 1.1 mm long single-shank 64-channel or double-shank 128-channel probe with tetrode configuration (Du et al. 2011; fabricated by the Masmanidis lab, UCLA, and assembled by the Litke lab, UCSC). The electrode was placed at an angle of 30-45 degrees to the cortical surface and inserted to a depth of 500-1000 *μ*m thebelow the cortical surface. A period of 30 minutes - 1 hour was allowed to pass before recording began. For each animal, the electrode was inserted only once.

### Visual Stimuli, Data Acquisition, and Analysis

Visual stimuli were presented as described previously (Niell and Stryker, 2008). Briefly, stimuli were generated in Matlab using Psychophysics Toolbox (Brainard, 1997; Pelli, 1997) and displayed with gamma correction on a monitor (Nanao Flexscan, 30 × 40 cm, 60 Hz refresh rate, 32 cd/m2 mean luminance) placed 25 cm from the mouse, subtending 60-75° of visual space. For Current Source Density (CSD) analysis, we presented a contrast-reversing square checkerboard (0.04 cpd, square-wave reversing at 0.5 Hz). To characterize neural responses with single unit recordings, we presented drifting sinusoidal gratings of 1.5 s duration at 100% contrast, with temporal frequency of 1 Hz, spatial frequency of 0.04 cycles/degree (cpd). We presented 12 evenly spaced directions in random order, interleaving a 0.5 s gray blank screen.

Movement signals from the optical mice were acquired in an event-driven mode at up to 300Hz, and integrated at 100 msec intervals. We then used these measurements to calculate the net physical displacement of the top surface of the ball. A mouse was said to be running on a single trial if his average speed for the first 500 ms of the trial fell above a threshold, found individually for each mouse (1-3 cm/s), depending on the noise levels of the mouse tracker. To make fair comparisons across behavior, we used an equal number of still and running trials in our analysis. This was done by finding the behavioral condition with the minimum number of trials (say N trials), and keeping only N trials (randomly chosen) from the other behavioral condition.

Data acquisition was performed using an Intan Technologies RHD2000-Series Amplifier Evaluation System, sampled at 20 kHz; recording was triggerec by a TTL pulse at the moment visual stimulation began.

### Single-neuron analysis

To find single-unit activity, the extracellular signal was filtered from 700 to 7 kHz, and spiking events were detected by voltage threshold crossing. Single units were identified using Vision Software (Litke lab, UCSF (Litke et al., 2004)) Typical recordings yielded 35-73 single units across the electrode. Neurons whose firing rates were unstable across the recoding session, characterized by a change of 75% in mean firing rate from the first third to the last third of the session, were excluded from further analysis. Units were classified as narrow- (putative inhibitory) or broad-spiking (putative excitatory) based on the shape of their average waveforms which were clustered into two groups using k-means on the first two principal components of waveform shape. Single-trial responses to visual stimuli were characterized as the number of spikes evoked during the first 500 ms after stimulus onset.

#### Cortical layer

Cell layer was estimated by performing current source density analysis (CSD) on data collected during presentations of contrast reversing square checkerboard. Raw data sampled at 20 kHz was first bandpass filtered between 1 and 300 Hz and then averaged across all 1 s positive-phase presentations of the checkerboard. Data from channels at the same depth were averaged together within a shank of the electrode; two mice had recordings from two-shank electrodes. CSD for each channel, *C_i_* was computed from the average LFP traces, *P* (*t*) using Eqn. 1, four site spacing, *s*, equal to a distance of 100 *μ*m).

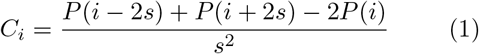

The borders between layers II/III-IV, IV-V, V-VI were identified by spatio-temporal patterns of sinks and sources in the CSD plot (Mitzdorf, 1985). The plot included in Figure 1c is of a 10x up-sampled CSD from mouse 1.

#### Cell tuning

Tuning curves for each neuron were found by taking a cell’s mean response across repetitions of a single visual stimulus. The change in spike count with locomotion was calculated as a function of mean spike count at rest:

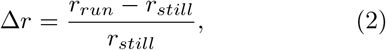

where *r* is the mean firing rate of the cell averaged across all stimulus conditions.

#### Additive and multiplicative modulation

Additive and multiplicative components of neural modulation were calculated by performing linear regression between the tuning curves fit separately to running and still data, treating the tuning curve in the still condition as the independent variable. The multiplicative coefficient obtained from the linear regression was taken to be the multiplicative component of modulation; the additive coefficient was further scaled by the mean firing rate across the tuning curves to compute the additive component of modulation. Modulation was assessed for significance using a re-sampling procedure, repeating the regression described above 1000 times on trials which were randomly assigned to the “still” or “run” categories. If the original regression coefficients fell outside of the 95% of the re-sampled distribution, they were considered significant.

#### Mutual Information

In the context of visually-evoked neural activity, a cell’s responses are considered informative if they are unexpected. For example, if a neuron in primary visual cortex consistently produces two spikes per second, the knowledge that the cell produced two spikes in response to a picture of a zebra does not provide any information. This notion can be formalized by a measure of information called the Shannon entropy (Shannon, 1948), the expected value of the information content of a particular variable, *H*(*X*) = *E_X_*[*I*(*x*)] = –∑*_x∈X_ p*(*x*)log_2_ *p*(*x*) computed here in units of bits. A neuron that has high variability of responses has high entropy, and is therefore said to be informative. The concept is further extended to mutual information, *I*(*X*_1_,*X*_2_), which quantifies how much information one variable contains about another. *I*(*X*_1_,*X*_2_) calculates the average reduction in uncertainty (entropy) about the first variable gained from knowing a particular instance of the second. Intuitively, a single response from a cell that is well tuned for visual grating orientation will leave little uncertainty as to the visual stimulus that evoked each response, whereas knowing the response of a poorly-tuned cell will result in little reduction in uncertainty. Mutual information between visual stimuli (*S*) and evoked single-neuron responses (*R*) is calculated as:

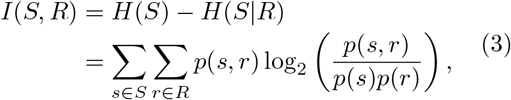

where *r* and s are particular instances from the set of neural responses (measured as spike counts) and stimuli (grating movement directions) respectively. The change in mutual information with locomotion was calculated as a function of *I*(*S*, *R*) at rest:

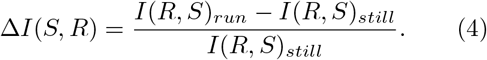

#### Stimulus-specific information

Stimulus-specific information, *SSI*(*s*), tells us how much information an average response carries about a particular visual stimulus, *s* (Butts 2003). Or, rephrased, it is the average reduction in uncertainty gained from one measurement of the response *r* ∈ *R* given a stimulus *s* ∈ *S*. The *SSI* of stimulus *s* is:

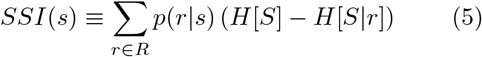

where *s*,*r*,*S*, and *R* are defined as above, and *H*[*S*] = –∑*_s∈S_ p*(*s*)log_2_ *p*(*s*)is the entropy of the visual stimuli and *H*[*S*|*r*] = –∑*_s∈S_ p*(*s*|*r*)log_2_ *p*(*s*|*r*) is the entropy of the visual stimuli associated with a particular response.

### Population-based analysis

#### Decoding visual stimulus from single-trial population responses

Data trials were separated into equal numbers of running and still trials, randomly subsampling from each 25 times to get a distribution of decoding errors based on the data included. We trained a linear discriminant analysis classifier to classify single-trial neural responses, assuming independence between neurons (a diagnonal covariance matrix), using a leave-one-out approach to train and test classification separately for the data from each behavioral state (LDA-LOOCV). The classifier was trained and tested using MATLAB’s *fitcdiscr* and *predict* functions. To decode only grating orientation and not movement direction, we grouped stimuli moving 180° apart into the same class.

#### Decoding from trials with equal population spike counts

To determine if firing rates are the sole determinants of information encoded within a neural population, we compared decoding accuracy from trials in running and still conditions with equal population spike counts, the sum of spikes from all neurons on a single trial. Although the distribution of population spike counts overlapped between rest and running, high population spike counts were more common during running and low population spike counts were more common at rest. To compare the two, we constructed a dataset that retained higher-order structure between neural activity with the population, but had many samples of running and still trials with the same population spike count. This was accomplished by performing LDA-LOOCV on different subsets of neurons from the population: 1 to 70 neurons were randomly sub-sampled from the population, yielding single-trial population spike counts that ranged from 0 to 275. For each number of neurons (e.g. 1, 5, etc.), we sub-sampled with replacement 100 times from the population, yielding 100 combinations of neurons. Classifiers were trained separately on each sub-sample and for each behavioral state (running vs. rest).

#### Signal and noise correlations

Using single-trial spike counts from the first 500 ms after stimulus onset, we calculated Pearson correlation coefficients for each pair of neurons recorded from a single mouse, *ρ_tot_.* These coefficients were assumed to be the sum of signal and noise correlations. Signal correlations, *ρ_s_*, measure similarity of tuning curves between neurons and were calculated by shuffling neurons’s responses to each visual stimulus. Noise correlations, *ρ_n_*, measure similarities in neural spiking across presentations of the same visual stimulus, and were calculated by taking the difference between total and signal pairwise correlations (*ρ_n_* = *ρ_tot_* – *ρ_s_*).

#### Decorrelating neural responses

Neural responses were decorrelated by randomly shuffling single cell responses to a particular stimulus across trials, where each trial was an instance of a single stimulus. For example, assume that *X* is an *n × m* matrix of neural responses, where *n* is the number of trials during which stimulus 1 was presented, and m is the number of neurons that were recorded. Shuffling randomly moves around entries in each column, so a single row will end up with neural responses from separate instances of a stimulus, and preserves the mean response of the cell to each stimulus while removing any correlations between neurons in time.

#### Stimulus discriminability, d’

Stimulus discriminability was calculated by taking all pairs of neighboring visual stimuli (*θ* ± *π*/4, movement in both directions) and plotting the population responses to each pair in n-dimensional space, where *n* is the number of neurons recorded from each mouse. We found the mean response for each stimulus, and projected each cloud of responses onto the vector between the two means. A d’ was calculated as:

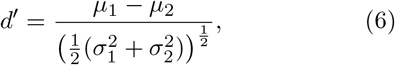

where *μ*_1_ and *μ*_2_ were the means of the projected data for each stimulus, and σ_1_^2^ and σ_1_^2^ were the variances. Discriminability was computed separately for each behavioral state.

## 3 Results

### Locomotion-induced modulation of evoked visual activity

We made stable simultaneous extracellular recordings from 36-73 single neurons in the primary visual cortex of each of eight awake, head-fixed mice that were presented with moving gratings in the monocular visual field contralateral to the recording site. Mice were free to run or stand stationary on a spherical treadmill floating on an air stream (Figure 1a) while their movements were monitored. Neuron spike times were extracted from raw data traces and sorted using custom software (Vision Software, Litke lab, UCSC). Experimental trials consisted of 1.5 s presentation of a moving grating following by 0.5 s of gray screen. Gratings could take one of 12 movement directions (two directions for each of 6 orientations), evenly spaced between 0 and 360 degrees. We characterized a neuron’s single-trial response to a visual stimulus by counting the number of spikes evoked in the first 500 ms after stimulus onset, labeling each trial as a “Run trial” or a “Still trial” based on the mouse’s average running speed during that period. Separate tuning curves of response as a function of grating movement direction for the Run and Still trials were calculated for each cell.

### Single cell evoked responses are more informative during locomotion

Average firing rates and mutual information (*I*(*S*, *R*), Eqn. 3) for single neurons were computed separately for each behavioral state, using a total of 240 to 961 trials (20 to 78 per stimulus), depending on the mouse. Fractional changes in firing rates and mutual information were calculated by dividing the change from rest to locomotion and normalizing it by the average value at rest. Locomotion strengthened average single-cell responses to the stimuli (mean increase of 62 ±93%, p = 1E-47, Wilcoxon signed-rank test; Figure 1b). The cortical layer in which a cell was located, found using current-source density analysis (Figure 1c), was related to the magnitude of a cell’s evoked responses, with cells in layers II/III and VI having the lowest evoked spike counts and those in layers IV and V having the highest (Figure 1d). The fractional changes in spike count were inversely proportional to the magnitude of evoked spike rates, but only the difference between layers II/III and layer V was significant (Figure 1e, Equation 2).

**Figure 1:**
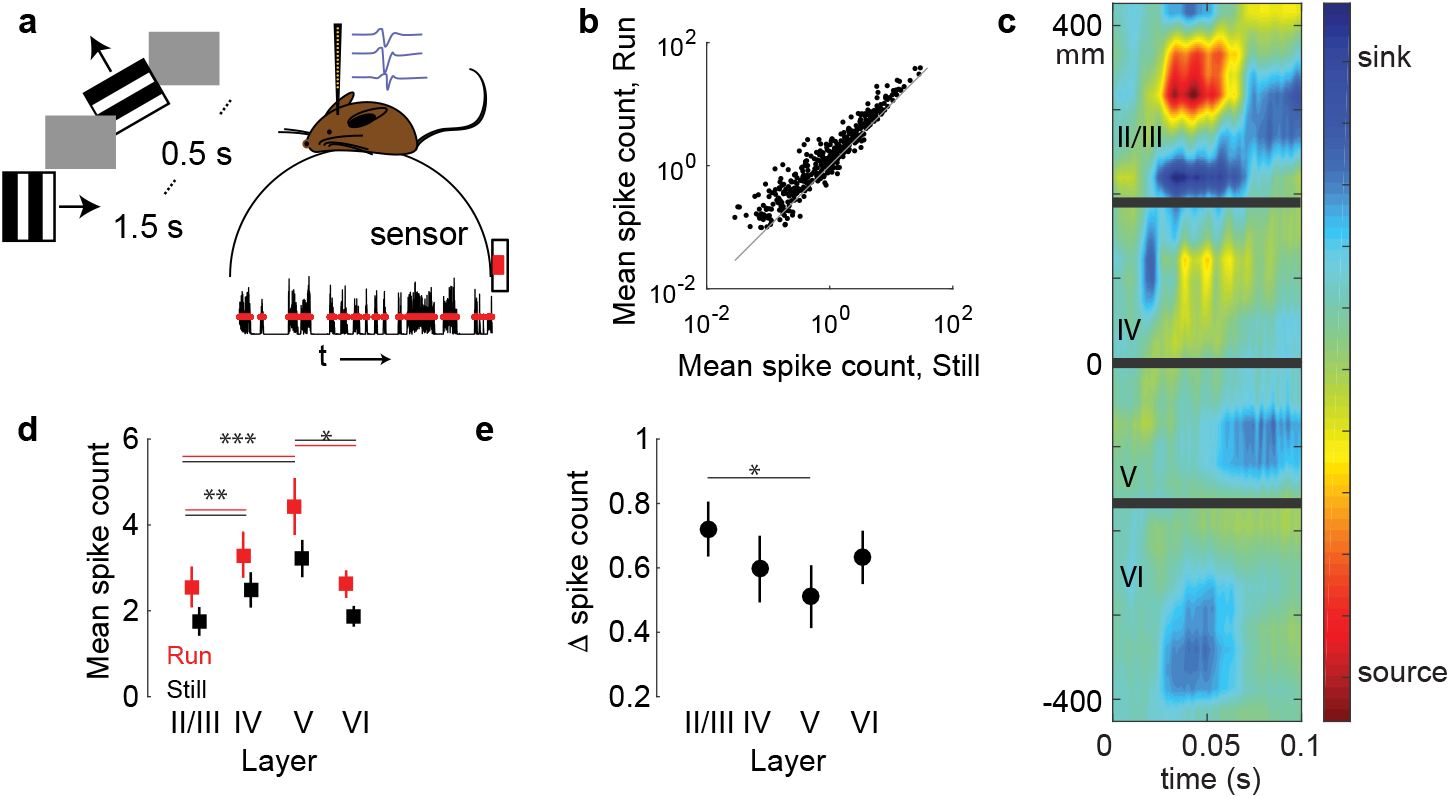
Cortical state change during locomotion. **a)** The activity of 35 to 73 single neurons was recorded from the primary visual cortex of freely-moving mice. Moving gratings (6 orientations, each moving in one of two possible directions) were displayed for 1.5s in the visual field contra-lateral to the recording site. Mouse movement was tracked over the course of the experiment. **b)** Evoked mean spike count, averaged over all visual stimuli, across behavioral conditions (N = 409 cells in 8 mice, p = 1E-47, Wilcoxon signed-rank test). Gray line is unity. **c)** Current-source density plot for one mouse overlaid with inferred laminar boundaries (thick gray lines). Distances at left refer to electrode location relative to the center of the array. **d)** Mean spike counts of cells by layer, averaged across all stimulus presentations. Layer II: N = 112, Layer IV: N = 90, Layer V: N = 84, Layer VI: N = 123. Error bars are bootstrapped estimates of standard error. * indicates p < 0.05, ** indicates p < 0.01, * * * indicates p < 1.3E-5, Wilcoxon rank-sum test. P-values were corrected for multiple comparisons using the HolmBonferroni method. **e)** Change in mean spike count during running as a fraction of mean spike count at rest, using data from **d**. Numbers of samples as in **d**. Error bars are bootstrapped estimates of standard error. * indicates p < 0.05, Wilcoxon rank-sum test. P-values were corrected for multiple comparisons using the HolmBonferroni method.

Running also increased mutual information between the responses and the set of visual stimuli, including the information encoded by a single spike (mean gain of 47 ±72%, p = 8E-31, Wilcoxon signed rank test; Figure 2a-b), calculated by diving *I*(*S*, *R*) by the mean spike rate of the cell, but the results across individual mice were more variable. While cells from all mice had significant shifts in *I*(*S*, *R*) per spike (p < 0.02, Wilcoxon signed rank test, Bonferroni correction for multiple comparisons), three of the eight mice actually had significant decreases in mutual information per spike with locomotion. Overall, 314 of 409 cells across all layers had increased *I*(*S*, *R*) with locomotion. The lower layers had the highest mutual information in both behavioral conditions (Figure 2c), possibly driven by high firing rates in layer V and the presence of a population of well-tuned cells in layer VI (Velez-Fort et al., 2014), but layers II/III had the largest fractional increase in mutual information during locomotion (Figure 2d). We next calculated a measure of stimulus-specific information (*SSI*(*s*), as defined in Equation 5) to determine if a cell’s information was directly proportional to its firing rate or if it depended on factors such as response variability. The amount of information a cell carried about each stimulus increased with the natural logarithm of mean spike count (Figure 2e), as is predicted by a Poisson encoder (see Appendix in (Ringach et al., 2002)).

As has been reported previously (Niell et al. 2010, Ayaz et al. 2013, Saleem et al. 2013, Fu et al. 2014, Lee et al. 2014, Erisken et al. 2014,Mineault et al. 2016), the responses of V1 neurons were modulated by locomotion. Response modulation consisted of both additive and multiplicative components (Figure 2f-h), which were computed by linearly regressing a neuron’s tuning curve during locomotion against its tuning curve at rest. The multiplicative coefficient obtained from linear regression was taken to be the multiplicative component of modulation; the additive coefficient was further scaled by mean firing rate across the tuning curves to compute the additive component of modulation.

**Figure 2:**
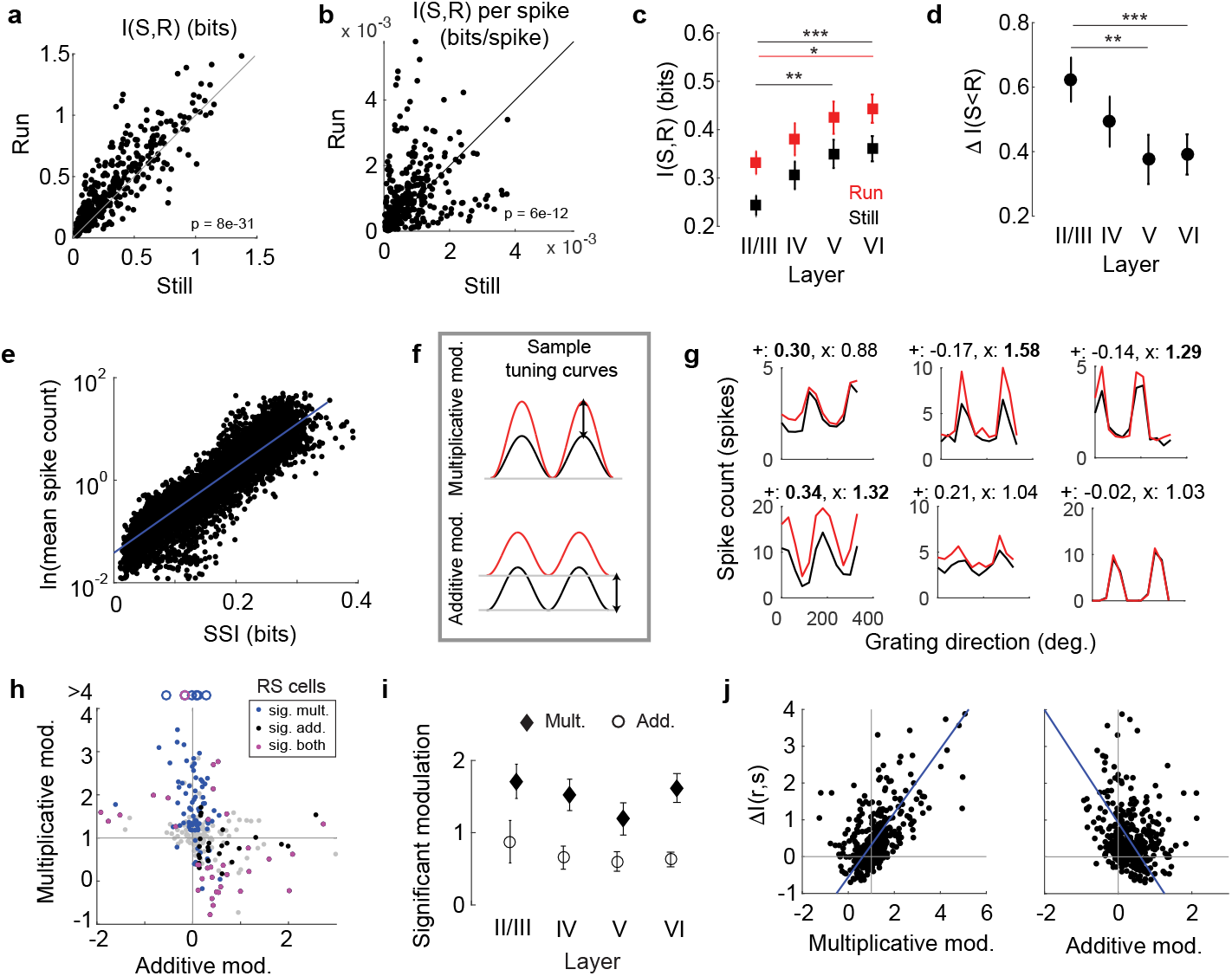
Cortical state affects single-neuron activity. **a.** Single cell mutual information, *I*(*S*, *R*), during running and rest (N = 409, p = 8E-31, Wilcoxon signed-rank test). Gray line indicates unity. **b.** *I*(*S*, *R*) per spike. (N = 409, p = 6E-12, Wilcoxon signed-rank test). Gray line indicates unity. **c.** Average behaviorally dependent *I*(*S*, *R*), within each cortical layer. Layer II: N = 112, Layer IV: N = 90, Layer V: N = 84, Layer VI: N = 123. Error bars are bootstrapped estimates of standard error. * indicates p = 0.012, ** indicates p = 0.002, * * * indicates p = 0.0007, Wilcoxon rank-sum test. P-values were corrected for multiple comparisons using the HolmBonferroni method. **d.** Fractional change in mutual information, Δ*I*(*S*, *R*), within each cortical layer during running. Error bars are bootstrapped estimates of standard error. ** indicates p = 0.002, * * * p < 0.0005, Wilcoxon rank-sum test. P-values were corrected for multiple comparisons using the HolmBonferroni method. **e.** Relationship between average spike count and stimulus-specific information, *SSI*(*s*). Each point is the SSI of a single cell to a particular grating movement direction (N = 4908). Blue line is fit of linear regression (R^2^ = 0.85, p = 0). **f.** Schematic of multiplicative (top) and additive (bottom) tuning curve shifts from rest (black) to locomotion (red). **g.** Sample single-cell tuning curves for evoked responses at rest (black) and during locomotion (red), with values of additive and multiplicative modulation printed above each. Bold indicates significant modulation. **h.** Relationship between additive and multiplicative components of modulation for each cell (N = 409, *ρ* = −0.315, p = 1.2E-7). Gray points are cells with no significant modulation, colors indicate significant modulation for multiplicative (blue), additive (black) or both (magenta) components. Gray represent null hypotheses that no modulation occurs. Open circles indicate data points outside of plot range. **i.** Average modulation across cortical layers during locomotion for cells that are significantly modulated. Error bars are bootstrapped estimates of standard error. **j.** Δ*I*(*S*, *R*) as a function of multiplicative (left; *ρ* = 0.58, p = 2.4E-38) and additive (right; *ρ* = −0.16, p =0.001) components of modulation (N = 409). Lines as described in **e**.

Across eight mice, 38% of neurons had significant multiplicative modulation (154/409, average of 1.5 ±1.3), 27% of neurons had significant additive modulation (110/409, average of 0.8 ±1), and 13% of neurons had both (54/409). Significance was computed using a resampling procedure (see Methods: Single-neuron analysis: Additive and multiplicative modulation). Average additive and multiplicative components varied across cortical layers, with layers II/III having the greatest multiplicative modulation and layer V the least (Figure 2i), although the difference was no longer significant after correcting for multiple comparisons. The change in the mutual information of cells with behavior was predicted by the multiplicative component of modulation, and was weakly inversely related to the additive component of modulation (*ρ* = 0.58, p = 2.4E-38 and *ρ* = −0.16, p = 0.001 respectively; Figure 2j). Although this change was necessarily driven by cells that were modulated, many of the most informative cells in the population were not modulated, and there was no significant relationship between the degree of additive or multiplicative modulation and mutual information in either behavioral state (p = 0.17 and p = 0.36 respectively, F-test of significance in regression).

### Populations of neurons encode more information about visual stimuli during locomotion

As single cell responses shift to encode more information about visual stimuli during locomotion, we might expect that the population as a whole would follow suit. Computing the mutual information between a neural population’s evoked responses and a visual stimulus would describe how well a population of neurons represents a visual stimulus; however, an accurate calculation of this value would require a vast number of trials. Instead, mutual information was estimated indirectly, by training a linear decoder (Linear Discriminant Analysis) on the data and asking how well visual stimuli could be predicted for single trials excluded from the training set. The classifier is linear, and makes several assumptions, including that evoked responses are independent across neurons and that they have a gaussian distribution. By comparing the accuracy with which single-trial responses could be classified, this technique allows comparison of how informative neural responses are about both the orientation and direction of the moving gratings during rest and locomotion.

Single-trial neural responses were classified more accurately during locomotion, both for the direction of grating movement (32% decrease in error, p = 3E-19, Wilcoxon sign test) and for grating orientation (44% decrease in error, p = 1E-18, Wilcoxon rank sum test; Figure 3a). Grating orientation was classified with higher accuracy than movement direction in both behavioral states, and the fractional improvement in its classification during locomotion was larger (44% vs. 32% decrease in error). The cells that were driving the change in information during running were not localized within a particular cortical layer: repeating the decoding analysis separately including only cells in layers II/II, IV, V, and VI yielded similar, significant changes in classification accuracy for each. (Figure 3a).

**Figure 3:**
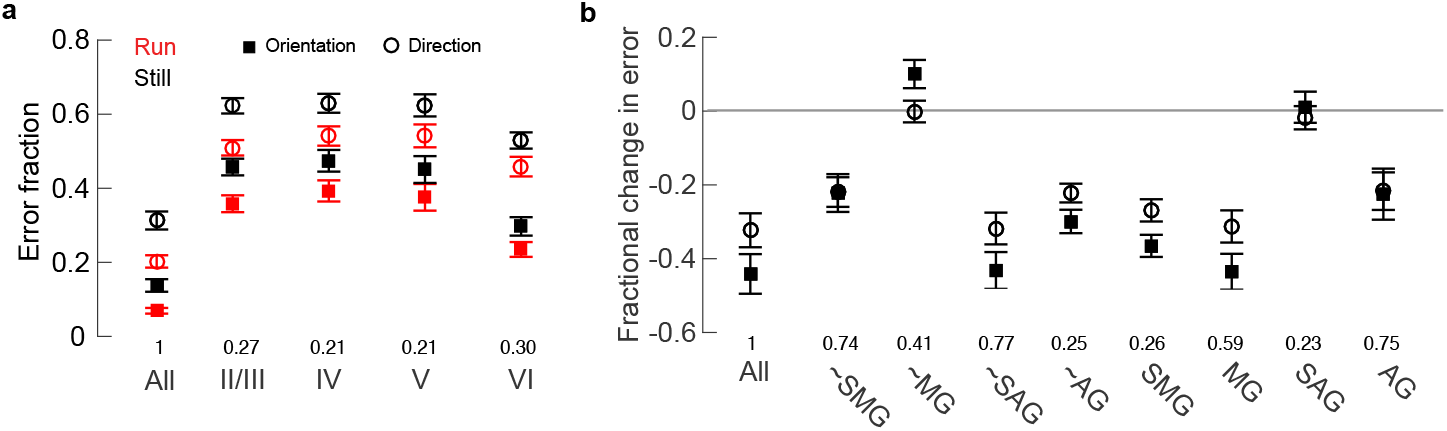
Classification of single-trial neural responses recorded during locomotion is more accurate than of those recorded at rest. **a)** Error in LOOCV-LDA classification of visual stimulus movement direction and orientation, as a population (All), and within particular layers. Numbers above layer labels denote the fraction of the total population included in the decoding. Error bars are bootstrapped estimates of standard error. **b)** Fractional change in decoding error with behavior (Error_*run*_ - Error_*still*_ / Error_*still*_). More negative values indicate greater improvement during locomotion. All: all cells, SMG: significant multiplicative gain > 1, SAG: significant additive gain > 0, MG: multiplicative gain > 1, AG: additive gain > 0.’∼’indicates the entire population excluding the category specified. Numbers above layer labels denote the fraction of the total population included. Error bars are bootstrapped estimates of standard error. Horizontal gray line indicates no change.

Next, to determine whether a particular subset of cells were most informative, we performed classification either by using only responses from that subset of cells or by using all cells in the population but that subset. Groups of interest included cells that had significant multiplicative gain greater than one (SMG), significant additive gain greater than zero (SAG), multiplicative gain greater than one (MG), and additive gain greater than zero (AG). Excluding cells with multiplicative gain eliminated the difference in classification error between rest and locomotion (Figure 3b.), revealing that these cells contributed most to this effect. When we excluded only SMG cells, the result was similar but less dramatic, presumably because many cells in which modulation did not reach significance by our criteria were actually modulated on most trials, leading to a difference in encoding accuracy across behavioral states. On their own, SMG and MG cells became far more informative during locomotion, statistically matching the fractional change in error observed when using the entire population of cells. In contrast, cells with significant additive modulation alone had no gain in information during running.

### Firing rates contribute to, but are not necessary for increased information content in a population

In the cortical state produced by locomotion, the information about the visual stimulus increases along with the visual responses of most neurons. Does the extra information available during locomotion result solely from the increase in neuronal firing rates, or does it also involve a change in the pattern of stimulus-evoked neural responses? Locomotion leads to higher population spike counts (the sum of spikes from all recorded neurons) on average, but the distributions of population spike counts during locomotion and rest have some overlap (Figure 4a). Comparing decoding accuracy in the two states for trials with equal population spike counts would preserve any higher order structure that might distinguish them, and would reveal whether information is exclusively determined by population spike counts. However, as the fraction of trials that directly overlapped is small, we generated a larger dataset with overlapping spike count by sub-sampling neurons from the population (see Methods: Population-based analysis: Decoding from trials with equal population spike counts.) When few neurons were sampled, the population spike count was forced to be low, and when many were sampled, it was high. Therefore, decoding accuracy was ultimately compared for equal population spike counts during rest and locomotion by including fewer cells in the locomotion classifier than in the rest classifier. LOOCV-LDA was performed separately for data collected during rest and during locomotion, after which the results from all mice were pooled together to generate average decoding error as a function of population spike count for each behavioral state. Classification error decreased with increasing spike count in both states, but the errors were lower for running trials than for still trials, even for equal population spike counts (Figure 4b), and particularly so at high population spike counts. These findings held both when classifying grating movement direction (left) and orientation (right). Thus, not just the amount but also the pattern of activity across the population is important for the accurately encoding visual stimuli, and locomotion shifts the population into a more informative state

**Figure 4:**
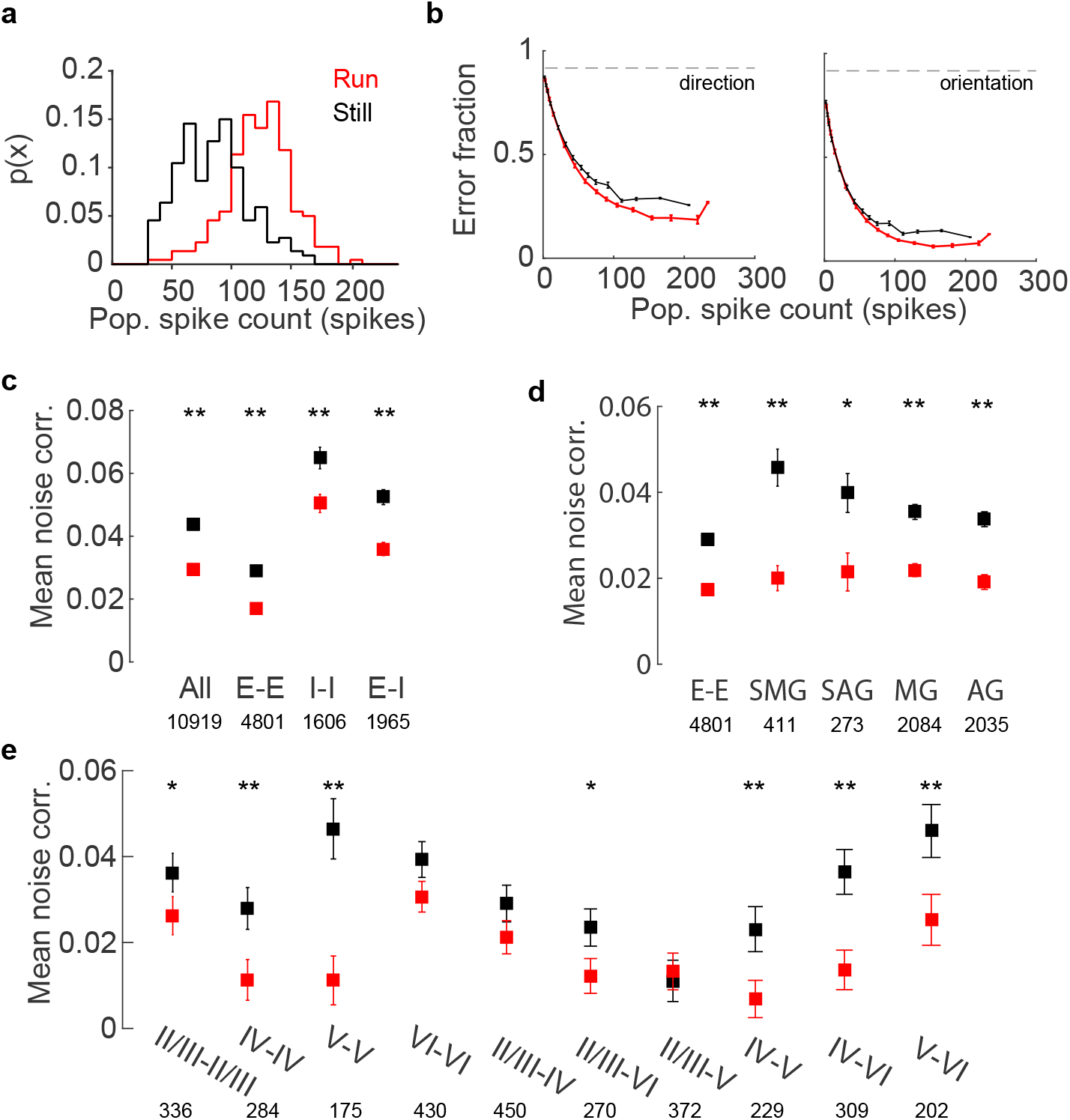
Noise correlations influence population representation of visual stimuli. **a)** The distribution of population spike counts, the sum of spikes from all neurons on a single trial, overlap in the running and rest conditions. **b)** Classification error for grating movement direction (left) and orientation (right) as a function of population spike count. Error bars are bootstrapped estimates of standard error. Dashed gray line denote chance levels of performance. **c)** Stimulus-independent (noise) pairwise correlations shift with behavior. Error bars are bootstrapped estimates of standard error. All: all pairs of cells, E-E: pairs of putative excitatory cells, I-I: pairs of putative inhibitory cells, E-I: pairs of one putative excitatory and inhibitory cells. Values below layer labels are number of pairs included in analysis. ** indicates significant change during running, p < 1E-5, Wilcoxon signed-rank test. **d)** Noise correlations between excitatory cells by cell modulation. Error bars are bootstrapped standard error of the mean. All: all pairs of excitatory cells, SMG: significant multiplicative gain > 1, SAG: significant additive gain > 0, MG: multiplicative gain > 1, AG: additive gain > 0. * and ** indicate significant change during running, p < 1E-3 and p < 1E-6 respectively, Wilcoxon signed-rank test. Values below labels are number of pairs included in analysis. **e)** Noise correlations between excitatory cells within a single layer and across layers. Error bars are bootstrapped estimates of standard error. Values below layer labels are number of pairs included in analysis. For significant changes during running, * indicates p < 0.02 and ** indicates p < 0.005, Wilcoxon signed-rank test.

### Locomotion decreases noise correlations

Stimulus discriminabilty, the extent to which visually-evoked neural responses differentiate visual stimuli, can be magnified or diminished by correlations between neurons (Cohen and Maunsell, 2009; Moreno-Bote et al., 2014). Correlated activity among neurons consists of two components: stimulus-dependent (signal) correlations that measure similarities in cell tuning, and correlated trial-to-trial fluctuations in response strength that are stimulus-independent (noise correlations). A reduction in pairwise noise correlations, as is observed during attention and locomotion via arousal (Cohen and Maunsell, 2009; Erisken et al., 2014; Vinck et al., 2015), could explain the improvement in classification that we observe during locomotion. Therefore we calculated Pearson pairwise correlations from trial-by-trial spike counts for each pair of neurons, separately for running and still conditions, and parsed these values into signal and noise correlations.

Locomotion had only a minor effect on average signal pairwise correlations (mean decrease of 0.003, p = 8E-7, Wilcoxon signed-rank test), but it substantially reduced mean noise correlations between all neurons (mean decrease of 0.014, p = 2.2E-50, Wilcoxon signed-rank test; Figure 4c). Noise correlations between putative excitatory-excitatory, inhibitory-inhibitory, and excitatory-inhibitory pairs significantly decreased, though pairs of inhibitory cells tended to have high, positive noise correlation in both behavioral states (Figure 4c). Pairs of putative excitatory cells with significant modulation were most decorrelated during locomotion (Figure 4d). Furthermore, excitatory cells across all cortical layers were decorrelated during running (Figure 4e). Layer V cells had the highest levels of noise correlations at rest, and were most decorrelated during running (mean decrease of 0.035, p = 3E-7, Wilcoxon signed-rank rest), followed by layer IV cells (mean decrease of 0.017, p = 0.002, Wilcoxon signed-rank test). The upper layers, layer II/III cells, were only moderately decorrelated during running (mean decrease of 0.01, p = 0.02, Wilcoxon signed-rank test), and layer VI cells were not significantly decorrelated (p = 0.17, Wilcoxon signed-rank rest). Furthermore, across layers, pairs of cells in layers II/III-IV, IV-V, IV-VI, and V-VI were decorrelated during locomotion.

### Increased firing rates and decorrelation improve stimulus discriminability

As noise correlations can either aid or hinder neural encoding (Averbeck et al., 2006; Ruff and Cohen, 2014; Moreno-Bote et al., 2014), the effect of reduced noise correlations on the population representation of visual stimuli is not obvious. However, it can be assessed indirectly by comparing the discriminability of population representations of two similarly oriented gratings, e.g. 0° and 30°, when single trial responses are decorrelated by shuffling (see Methods: Population-based analysis: Decorrelating neural responses) to when correlations are preserved. The discriminability of a pair of stimuli can be measured by calculating *d’* of response distributions (the difference in their mean responses divided by the root mean square of their standard deviations;Cohen et al. 2009; Figure 5a; see Methods: Population-based analysis: Stimulus discriminability, d’). We applied this analysis to the neural representations of oriented gratings, calculating *d’* for neighboring pairs of grating movement directions, *θ* with 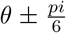 and *θ* with 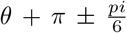. As expected, *d’* values were higher for pairs of stimulus representations observed during locomotion than during rest, implying that visual stimuli should be better separated in the neural response space during locomotion (average 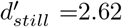, 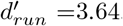, mean increase of 47%, p = 3E-17, Wilcoxon signed-rank test; Figure 5b-c).

**Figure 5:**
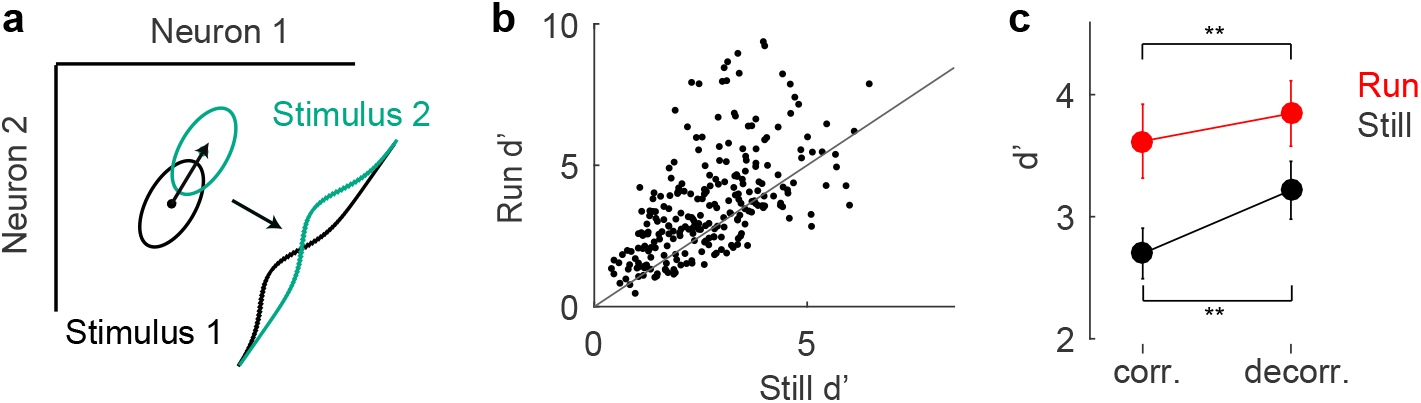
Stimulus discriminability depends on firing rates and noise correlations. **a.** Schematic calculation of d’ measure. Ovals are distribution of responses of two neurons to two visual stimuli. Black arrow between response distributions illustrates difference vector upon which responses are projected, yielding distributions drawn in lower right. Values for d’ are calculated from these overlapping distributions using Eqn. 6. **b.** Discriminability of grating movement direction, calculated on pairs of neighboring stimuli across behavioral state, p = 3E-17, Wilcoxon signed-rank test. Black points: d’ for a pair of stimuli (n=248; 8 mice, 31 per mouse). Gray line: unity. **c.** Decorrelation reduces change in d’ with behavior, and increases overall d’ values. Mean improvement in d’ with correlated data: 47%, p = 3E-17, Wilcoxon signed-rank test. Mean improvement in d’ with decorrelated data: 31%, p = 1E-12, Wilcoxon signed-rank test. Error bars are bootstrapped confidence intervals of the mean. ** indicate p < 5E-8, for difference between correlated and decorrelated d’ values.

To isolate the separate effects of increased spiking and decorrelation, we examined the change in d’ when one factor was held constant. First, to assess the effect of an increase in spike count when noise correlations were held fixed, we calculated d’ for populations whose responses had been decorrelated by shuffling. These shuffled populations lack any noise correlations, so comparing d’ across behavior reveals only the effect of increasing spike counts. Decorrelating responses in this way substantially reduced the effect of behavior on discriminability but did not eliminate it (average 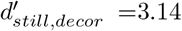, average 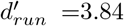, mean increase of 23%, p = 6E-16, Wilcoxon signed-rank test; Figure 5c). Therefore locomotion improves stimulus discriminability not only by increasing the distance between the mean responses to different stimuli through increases in firing rates, but also by reducing variability in responses through decorrelating responses.

### Time course of Information

When the mouse is at rest, the brain has unlimited time to integrate information from the stable visual scene, but during locomotion the visual system must encode the scene swiftly. In both cases, visually-evoked responses are dynamic, beginning with a sharp onset around 50 ms after stimulus presentation, then falling to a stable, elevated rate for the remainder of the stimulus presentation. How much information about the visual stimulus do cells contain at different points over the course of the response, and at what relative stimulus durations are the information content of these two states equivalent (e.g. at what stimulus duration will decoding from responses at rest yield the same decoding accuracy as decoding from the first 100 ms during locomotion)?

To determine if single cell responses were more informative during locomotion throughout the duration of the evoked response, we computed mutual information in ten millisecond bins. Average singlecell *I*(*S,R*) closely followed the time course of spike rates (not shown), and *I*(*S*, *R*) during locomotion was higher than that at rest for the entirety of the evoked neural response (≈50-500 ms; Figure 6a). Therefore, in single cells, cortical state change during locomotion confers a persistent, not transient, advantage in representing visual stimuli. We next compared the amount of information in the entire neural population at different time points, using LDA-LOOCV to estimate grating direction and orientation from spike counts during four sequential 100 ms periods, beginning with the time of response onset, some 50 ms after the stimulus was first presented. Consistent with single-cell mutual information, the population of neurons was most informative during the first 100 ms after neural response onset, with smaller decoding errors than during subsequent 100-ms periods (Figure 6b). Unsurprisingly, using data from the entire 500 msec period was superior to even the most informative 100 ms period, revealing that information is gained with longer periods of integration, regardless of cortical state.

**Figure 6:**
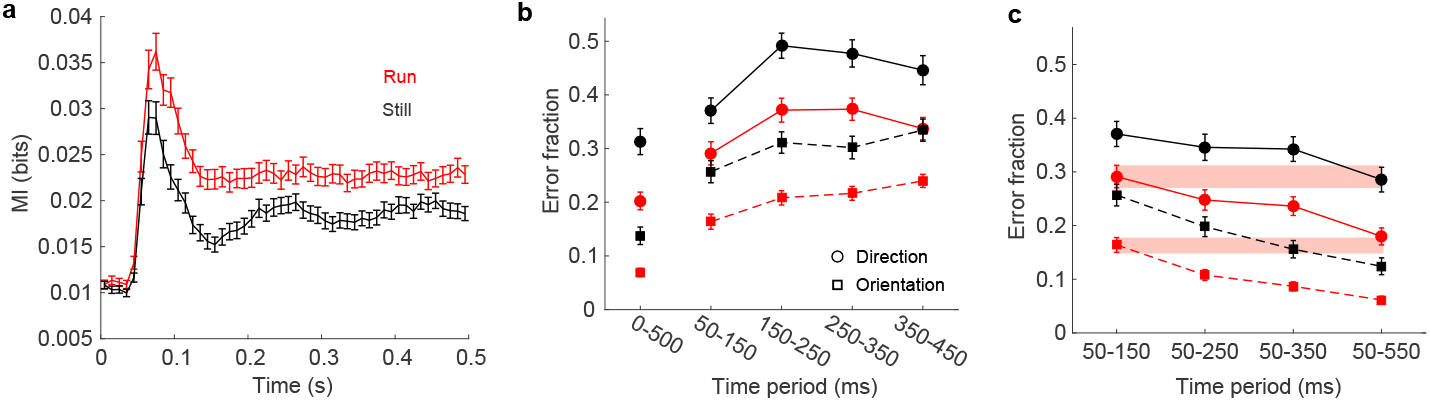
Stimulus information as a function of time. **a.** Mutual information between cell spiking and stimulus per 10 ms time bin, averaged across all recorded cells. Error bars are bootstrapped estimates of standard error. **b.** Classification error over sequential 100 ms time periods after stimulus onset. Error bars are bootstrapped estimates of 95% confidence intervals. **c.** Classification error over various time ranges after stimulus onset. Red is locomotion; black is still. Error bars are bootstrapped estimates of 95% confidence intervals. Shaded bars represent 95% confidence intervals of the mean during the first 100 ms after neural response onset.

In order to find a point of equivalence between decoding errors in the two behavioral states, we compared classification errors on population responses over a range of stimulus durations: 50-150 ms, 50-250, 50-350 ms, and 50-550 ms (Figure 6c). Classification accuracy achieved using spike counts from the first 100 ms of run trials was equal to that using spike counts from the first 300 ms (for stimulus orientation) or 500 ms (for stimulus movement direction) of still trials. It therefore takes vastly different times for the two states to yield similar net information.

### Are cortical states binary?

The information encoded in the population grows with spike count, but single-cell spike counts are only slightly modulated by the running speed. Indeed, in only 72 of 409 cells was more than 1% of the variance in spiking explained by linearly regressing spike counts against run speeds. Furthermore, residual spike counts, calculated by subtracting each cells’s mean response to a visual stimuli from its evoked response, were only weakly related to running speed (Figure 6a). Similar, but qualitative, observations were reported in excitatory neurons in V1 (Niell and Stryker, 2010) and in the inhibitory neurons thought to convey information about locomotion to V1 (Fu et al., 2014). Then, to what extent is population-level information proportional to mouse running speed?

To answer this question, we repeated the LDA-LOOCV analysis on just running trials and examined the relationship between run speed, population spike counts, and classification error. As shown above, single-trial population spike counts were predictive of classification error (Fig. 4b). However, population spike count was only weakly predicted by a linear function of run speed or the natural logarithm of run speed (Figure 7b). Instead, more than 99% of the variability single-trial population spike counts was left unexplained, even though the relationship between variables was significant in all of the mice. As forecast by the preceding results, run speed was not significantly predictive of average classification error in individual mice (Figure 7c). Classification error saturated with increases in running speed over 1-2 cm/sec when we considered responses during the first 500 msec after stimulus onset (Figure 7c). In only 3 of the 8 mice did error decrease significantly (p ≤ 0.04) with running speed, suggesting that, at least in most mice, the effect of locomotion on stimulus encoding is more nearly binary than graded.

**Figure 7:**
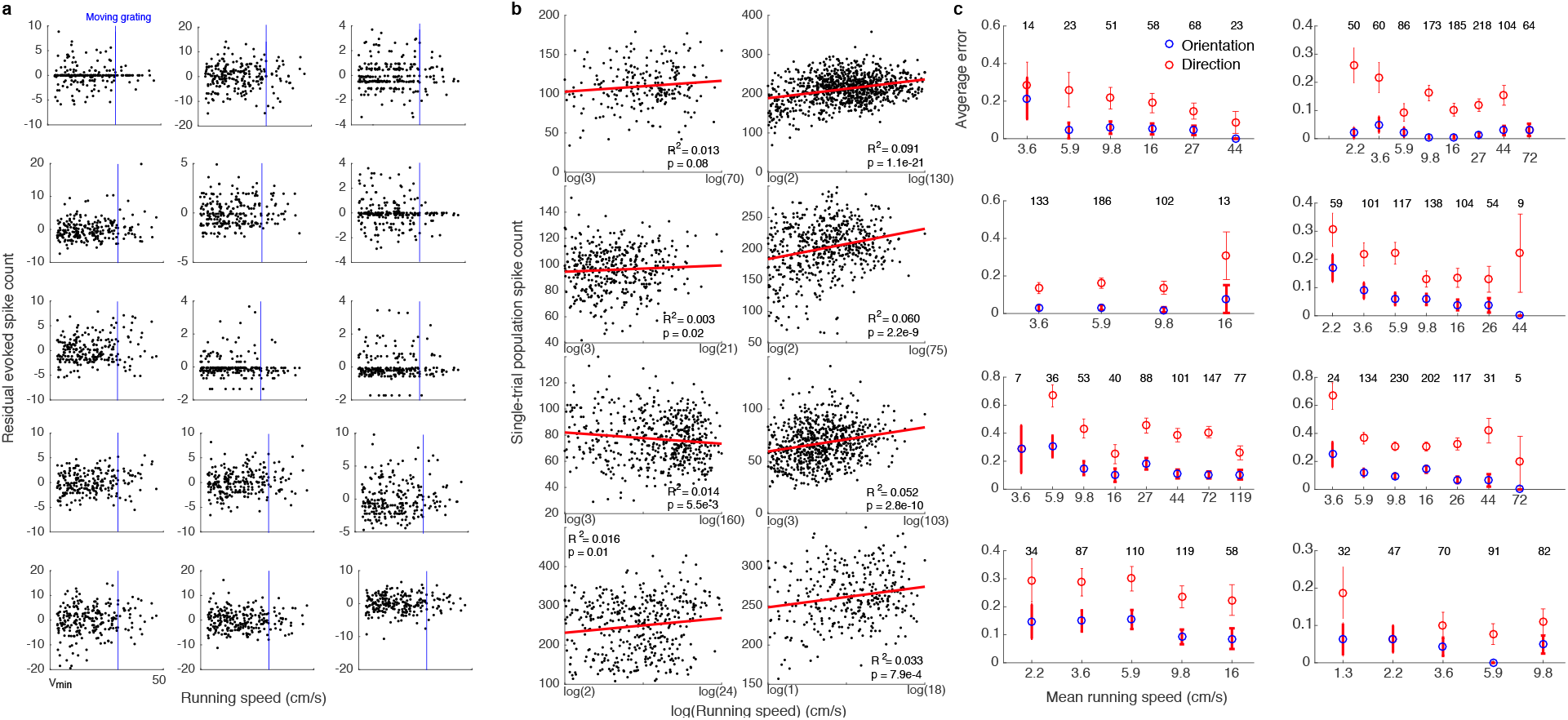
Relationship between running speed and spike counts, population responses, and classification error. **a.** Residual spike counts as a function of run speed for fifteen sample cells after mean visually-evoked responses were subtracted. Cells were chosen randomly from the population in a single mouse; responses shown are from run trials. Blue bar: speed of visual stimulus, 30 cm/s. **b.** Population spike counts as a function of the natural logarithm of mouse running speed on single running trials (black dots). Red lines, R^2^ values, and p-values indicate fit of linear regression. Panels are individual mice. **c.** Average LOOCV error with increasing mouse running speed for stimulus orientation (blue) and movement direction (red). Numbers of samples at each mean speed are listed at top of each panel.

## 4 Discussion

### Summary

Our data demonstrate that mouse V1 represents visual information with higher accuracy during locomotion than at rest, reducing the time required to correctly portray visual inputs. This is accomplished by a change in cortical state across the depth of the cortex that both increases firing rates of single cells and decorrelates non stimulus-related spiking among cells. Not only does the amount of information conveyed by V1 increase with locomotion, but, on average, the information conveyed by each spike within the population increases, although the process by which this is accomplished varies across cortical layers. Furthermore, the effect does not seem to be graded by movement speed and instead is closer to a binary switch in cortical state. Together, these changes should allow the mouse visual system to process the dynamic visual scenery experienced during running accurately and rapidly.

### Behavioral modulation of the neural code

Behaviorally-induced, rather than random, fluctuations in cortical state may have greater effects on population-wide encoding in V1. For example, in a recent report on monkey primary visual cortex (Arandia-Romero et al., 2016), spontaneous transitions from low to high population activity did not alter the total information available about grating orientation. Instead, it appeared that the gain in information from neurons that were multiplicatively modulated was offset by the loss from neurons that were additively modulated. In contrast, but in agreement with our present result, a study examining the effect of locomotion on neurons in layers II/III of mouse visual cortex found that grating orientation was easier to read from population activity during locomotion, and that the greatest gains were made for stimuli with high spatial frequency (Mineault et al., 2016). The present study additionally shows that decoding accuracy of both grating orientation and movement direction (for stimuli at a fixed spatial frequency) is enhanced for neurons in deeper layers of cortex, even though these neurons tend to have lower multiplicative gain values (Erisken et al., 2014) and a smaller fractional change in mutual information. These findings suggest that spontaneous shifts in population activity may have little significance, but behaviorally-elicited changes affect information transmission in the animal models studied.

### Specificity of results to cortical layers

Although locomotion increased the accuracy with which visual stimuli were decoded from evoked neural activity in every cortical layer (Figure 3a), these changes seem to have been driven by distinct mechanisms in each: cells in layers II/III underwent a large increase in mean firing rates relative to baseline and a small but significant decrease in noise correlations, cells in layer V had only a small increase in fractional firing rates but experienced a large decrease in noise correlations, and cells in layers IV and VI fell somewhere in-between, and probably result from some combination of the processes described below.

The increase in layer II/III firing rates has been explained by a disinhibitory circuit model, where cholinergic inputs from the basal forebrain excite VIP-positive interneurons that in turn inhibit somatostatin-positive interneurons (SST), effectively disinhibiting excitatory neurons in V1 (Fu et al., 2014). In contrast, layer V VIP cells are fewer (Lee et al., 2010) and morphologically distinct (Pronneke et al., 2015) from those in layers II/III, and they only weakly inhibit SST cells (Pfeffer et al., 2013). Therefore, in layer V, only a small change in firing rates can be expected during locomotion. Note, however, that contradictory reports of SST behavior in mouse V1 during locomotion (Polack et al., 2013; Fu et al., 2014; Reimer et al., 2014; Pakan et al., 2016) has led to the development of an alternative model of interneuron activity in layers II/III: VIP and SST cells are mutually inhibitory and their relative activity is dependent on the type of visual input available. The disinhibitory circuit described previously is presented as a sub-case that occurs when visual inputs are small, thus strongly exciting VIP cells but only weakly activating SST cells, leading to disinhibition. Large visual inputs, as were used in the experiments described here, robustly drive both cells types; however, as SST cells receive greater net input, they dominate and inhibit both VIP and pyramidal cells. It is not clear under such a model how pyramidal neurons increase firing rates during locomotion.

The second mechanism, a decrease in noise correlations during locomotion (Erisken et al., 2014; Vinck et al., 2015), is driven by heightened arousal (Reimer et al., 2014; Vinck et al., 2015). In general, pairwise noise correlations in pyramidal cells are thought to result from fluctuations in drive to neurons by non-sensory factors (Ecker et al., 2010; Goris et al., 2014; Reimer et al., 2014; McGinley et al., 2015; Vinck et al., 2015), which shift the magnitude of feedback inhibition to increase (less inhibition) or decrease (more inhibition) noise correlations (Stringer et al., 2016). For example, cholinergic projections from the basal forebrain can decorrelate neural population responses (Goard and Dan, 2009) by directly exciting SST neurons (Chen et al., 2015). If this circuit explains the shift in noise correlations with locomotion, layers that exhibit substantial reductions during locomotion should have SST cells as a significant portion of interneurons and should receive cholinergic inputs from the basal forebrain. Indeed, SST cells comprise just under half of all interneurons in layer V (Lee et al., 2010), where noise correlations were profoundly reduced during locomotion (Figure 4e), and the lower portion of this layer receives cholinergic inputs (Kitt et al., 1994). The relative balance of cholingeric inputs and interneuron distribution and connectivity may explain the differences observed in noise correlations across cortical layers. Overall, quick shifts in wakefulness of the animal could have inflated our estimates of noise correlations, both while mice are at rest and during locomotion (Reimer et al., 2014; McGinley et al., 2015; Vinck et al., 2015).

### Computational goal of cortical state change

Two additional explanations have been advanced for behaviorally-driven shifts in neural firing patterns. The first posits that neurons in layers II/III of mouse V1 are encoding sensory mismatch signals, the difference between expected and true visual flow given the mouse’s run speed (Keller et al., 2012), while the second suggests that neurons in V1 represent an integrated estimate of visual flow and running speed of the mouse (Saleem et al., 2013). They both suggest that motor information, perhaps efference copy, is transmitted to mouse V1, either to differentiate between self-generated and external visual flow or to help the mouse estimate his own movement speed. The object of this paper is not to refute either of these hypotheses, but to argue for an additional, third purpose for the modulation of neural firing rates in mouse V1 during locomotion. As both studies used a virtual reality environment to manipulate the relationship between visual flow and running speed, our results cannot be directly compared. However, these hypotheses make specific predictions, and we can ask if the explanations they pose towards elevated firing rates during locomotion can explain the pattern of results in the present study.

If neurons were encoding sensory mismatch, the most vigorous neural responses would be elicited when the difference between movement speed and visual speed were largest. Instead, we found that neurons, including those in layers II/III, had visually-evoked responses that were only weakly modulated by mouse running speed above 1-2 cm/sec (Figure 7a), and were not minimal at around 30 cm/s (the movement speed of the visual stimulus), contradicting the notion that sensory mismatch explains our results.

If neurons were integrating visual speed and locomotor speed, neural responses would be best explained by a function of both. As our data were generated using a fixed visual stimulus speed, we could only study the effect of running speed on neural responses in V1. As described above, the neural population became more informative at higher running speeds in only a minority of the mice in the present study, and only weakly so (Figure 7). Furthermore, neither at the level of single neurons nor at the level of population activity did spike count substantially rise with running speed, which is an important determinant of information content at both the single cell and population levels.

We propose that enhanced information processing, representation of sensory mismatch, and sensorimotor integration, may all be taking place simultaneously in V1. Perhaps motor input to V1, in the form of efference copy from sensorimotor areas, allows mice to differentiate between internally- and externally-generated visual flow, while cholinergic inputs from the basal forebrain modulate the gain of neuronal responses to improve information coding. A similar heterogeneity exists in primary somatosensory cortex of macaques, which has cells that primarily respond to sensory input, others that respond to motor signals, and others that are modulated by a combination of the two (London and Miller, 2013). We may expect a comparable mixture in mouse primary visual cortex.

## Acknowledgments

This work was supported by a Simons Collaboration on the Global Brain Grant to M.P.S, a Simons Collaboration on the Global Brain Fellowship to M.C.D., and National Eye Institute Grant R01 EY02874 to M.P.S., and a RPB Stein Innovation Award to M.P.S. We thank the Masmanidis lab at UCLA for electrode fabrication, the Litke lab at UCSC for electrode assembly and the Vision spike sorting software, and J.G. Makin and Y.J. Sun for thoughtful discussions and ideas.

